# Year-round quantification, structure and dynamics of epibacterial communities from diverse macroalgae reveal a persistent core microbiota and strong host specificities

**DOI:** 10.1101/2024.07.22.604553

**Authors:** Maéva Brunet, Nolwen Le Duff, Tristan Barbeyron, François Thomas

## Abstract

Macroalgae-bacteria interactions play pivotal ecological roles in coastal ecosystems. Previous characterization of surface microbiota from various macroalgae evidenced fluctuations based on host tissues, physicochemical and environmental parameters. However, the dynamics and degree of similarity of epibacterial communities colonizing phylogenetically distant algae from the same habitat are still elusive. We conducted a year-long monthly epimicrobiota sampling on five algal species inhabiting an English Channel rocky shore: *Laminaria digitata*, *Ascophyllum nodosum*, *Fucus serratus* (brown algae), *Palmaria palmata* (red alga) and *Ulva* sp. (green alga). To go beyond relative compositional data and estimate absolute variations in taxa abundance, we combined qPCR measurements of 16S rRNA gene copies with amplicon metabarcoding. A core microbiome composed of 10 genera was consistently found year-round on all algae. Notably, the abundant genus *Granulosicoccus* stood out for being the only one present in all samples and displayed an important microdiversity. Algal host emerged as the primary driver of epibacterial community composition, before seasonality, and bacterial taxa specifically associated with one or several algae were identified. Moreover, the impact of seasons on the epimicrobiota varied depending on algal tissues. Overall, this study provides an extensive characterization of the microbiota of intertidal macroalgae and enhances our understanding of algal-bacteria holobionts.

## Introduction

Macroalgae play crucial roles in marine ecosystems, contributing significantly to primary production, nutrient cycling, and habitat provision. Their surface harbors a dense, complex and highly dynamic biofilm of bacteria, archaea, fungi and viruses. Bacteria dominate the microbiota in abundance and diversity, representing up to 10^8^ cells.cm^-2^ (Egan *et al*., 2013; Martin *et al*., 2015), and form intricate interactions with macroalgae. These associations were reported to be either beneficial, detrimental or commensal (Burgunter-Delamare *et al*., 2024) and are at the basis of various essential processes in coastal oceans, including roles in nutrient cycling (Brunet *et al*., 2022) and in shaping host morphology, reproduction and defense against pathogens (Wichard *et al*., 2015; Singh and Reddy, 2016). The structure and composition of the microbiota of various brown (eg. Bengtsson et al., 2012; Florez et al., 2019; Burgunter-Delamare et al., 2023), red (Miranda *et al*., 2013; Zozaya-Valdés *et al*., 2017) and green (Burke *et al*., 2011; van der Loos *et al*., 2023) macroalgae has been largely investigated in the past decades using culture-dependent and -independent approaches, allowing for the identification of key members of the epiphytic microbiota. Macroalgae host specific bacterial communities that largely differ from those occurring in the surrounding seawater (Weigel and Pfister, 2019; Lu *et al*., 2023). Among bacteria associated with macroalgae, the bacterial phyla *Pseudomonadota* (mostly the classes *Alphaproteobacteria* and *Gammaproteobacteria*), *Bacteroidota*, *Verrucomicrobiota*, *Actinomycetota* and *Planctomycetota* are the most represented (Wahl *et al*., 2012; Hollants *et al*., 2013; Florez *et al*., 2017). At the genus level, the algal symbiont *Granulosicoccus* (*Gammaproteobacteria*) is commonly found as the most abundant taxon on diverse algal hosts (Singh and Reddy, 2014), frequently reported to account for more than 25% of the whole community on kelp tissue (Brunet *et al*., 2021; Lemay *et al*., 2021; Ramírez-Puebla *et al*., 2022; Weigel *et al*., 2022). It is now well established that bacterial community composition and density is shaped by biotic and abiotic factors and varies depending on biogeography, host tissue composition or seasonality (Lachnit *et al*., 2011; Egan *et al*., 2013; Lemay *et al*., 2018; Serebryakova *et al*., 2018; Weigel and Pfister, 2019; Paix *et al*., 2021; Ramírez-Puebla *et al*., 2022). Algae release a wide range of bioactive compounds with a broad spectrum of activities that participate in the control of bacterial colonization (Saha and Fink, 2022). It was reported that host taxonomy (Kuba *et al*., 2021; Chen *et al*., 2022), anatomy (Bengtsson *et al*., 2012; Weigel and Pfister, 2019; Lemay *et al*., 2021), genetics (Wood *et al*., 2022), health (Fernandes *et al*., 2012; Marzinelli *et al*., 2015; Burgunter-Delamare *et al*., 2023) or life cycle (Bengtsson *et al*., 2010; Glasl *et al*., 2021) influences its surface microbiota. Studies investigating seasonal variations in bacterial communities associated with healthy macroalgae have revealed dynamic patterns characterized by shifts in community composition and diversity (Lachnit *et al*., 2011; Burgunter-Delamare *et al*., 2023). These shifts can often be linked to seasonal changes in environmental factors such as temperature, light or nutrient availability (Mancuso *et al*., 2016; Florez *et al*., 2019), which influence the growth and metabolic activity of both macroalgae and associated bacteria. Additionally, seasonal variations in macroalgal physiology, such as growth rates and reproductive cycles, can indirectly impact bacterial communities by altering the tissues and exudate composition (Pandey *et al*., 2022).

Most of the works that characterized macroalgal-associated microbiota mainly focused on a single algal species. Only a few recent studies compared algae from different phyla and of distinct chemical composition revealing that algal phylum was a strong driver of epimicrobiota composition (Kuba *et al*., 2021) and identifying core taxa (Chen *et al*., 2022; Lu *et al*., 2023). Moreover, temporal dynamics of epiphytic bacterial communities depending on the algal host and type of tissue have been largely overlooked. In this study, the microbiota of three brown (*Laminaria digitata* (Hudson) J.V.Lamouroux, 1813, *Fucus serratus* Linnaeus, 1753 and *Ascophyllum nodosum* (Linnaeus) Le Jolis, 1863), one red (*Palmaria palmata* (Linnaeus) F.Weber & D.Mohr, 1805) and one green (*Ulva* sp. Linnaeus, 1753) macroalgal species predominant in the English Channel were investigated throughout a year. This selection of species covers a large phylogenetic range, and different intertidal zonation (*A. nodosum* is found on the upper-middle intertidal zone while *F. serratus*, *P. palmata* and *Ulva* sp. are in the lower intertidal zone and *L. digitata* in the subtidal zone (Evans, 1957; Burel *et al*., 2022)). The sole use of metabarcoding amplicon sequencing has emerged as the preferred method to investigate host-associated microbial communities. The resulting datasets are inherently compositional as they only allow analysis based on relative proportions of microbial taxa (Gloor *et al*., 2017). Therefore, an increase in abundance of one taxon can partly be caused by an equivalent decrease of another one, limiting the conclusions that can be drawn (Jian *et al*., 2020). Despite the statistical approaches applied to overcome misinterpretations of bacterial community structure (Gloor *et al*., 2017; Shelton *et al*., 2022), one of the most effective ways to accurately compare microbial communities across samples and to identify patterns that may be obscured when relying solely on relative abundance information, is to generate quantitative microbial abundance data by combining amplicon sequencing with the cell or gene copy density measurements. Several studies demonstrated an improved performance of quantitative approaches over compositional data (Jian *et al*., 2020; Lloréns-Rico *et al*., 2021) and this has been recently implemented to examine human (Vandeputte *et al*., 2017; Barlow *et al*., 2020), freshwater (Props *et al*., 2017) or soil (Camacho-Sanchez, 2023) microbiomes. Here we coupled 16S rRNA gene amplicon sequencing with quantitative PCR to accurately estimate the abundance of the different taxa in macroalgal epiphytic communities. We aimed to (i) characterize the shared and specific patterns of epiphytic bacterial communities associated with co-occurring macroalgae species and (ii) assess if seasonality equally impacts microbiomes from phylogenetically distant species.

## Experimental procedures

### Sampling

The collection of epibiota from algal specimens was previously described in (Brunet *et al*., 2023). Briefly, the surface microbiota of intertidal algae was sampled monthly between February 2020 and January 2021 (no sampling in April and May 2020 due to the Covid-19 pandemic). Microbiota were collected in triplicates using sterile flocked nylon swabs (Zymobiomics) on healthy specimens of the brown algae *Laminaria digitata* (Ldig, 0.5-1 m long), *Fucus serratus* (Fser) and *Ascophyllum nodosum* (Anod), the red alga *Palmaria palmata* (Ppal) and a green alga *Ulva* sp. (Ulva) at the Bloscon site (48°43’29.982’’ N, 03°58’8.27’’ W) in Roscoff (Brittany, France). Swabbed surface was standardized to 50 cm^2^. Three different regions of the kelp *L. digitata* were sampled: the basal meristem (young tissue, hereafter LdigB), the medium frond (ca. 20 cm away from the meristem, hereafter LdigM) and the old frond (the blade tip, hereafter LdigO). Only two replicates were retrieved for LdigB in February and Ulva in March. Upon collection, swabs were immediately immersed in DNA/RNA Shield reagent (ZymoBiomics) on ice and stored at -20 °C until DNA extraction. A total of 209 samples were collected (**SuppFile S1**), scattered into 70 different conditions (7 types of algal tissues at 10 different months).

### Collection of environmental data

A set of 15 environmental parameters measured at 1.3 km from the collection site (Estacade sampling station) between February 2020 and January 2021 was used for correlation analyses. It includes surface seawater temperature (T, °C), salinity (S), dissolved oxygen (O, ml*l^-1^), pH, ammonium (NH_4_, μM), nitrate (NO_3_, μM), nitrite (NO_2_, μM), phosphate (PHO_4_, μM), silicate (SiOH_4_, μM), particulate organic carbon (COP, μg*l^-1^) and nitrogen (NOP, μg*l^-1^), suspended matter (MES, mg*l^-1^), ^15^Nitrogen (DN15, °/_°°_) and ^13^Carbon isotopes (DC13, °/_°°_) and Chlorophyll a (CHLA, μg*l^-1^). Data were downloaded from the SOMLIT database (Service d’Observation en Milieu Littoral; http://www.somlit.fr, Cocquempot et al., 2019) on April 7th, 2022 and are listed in **SuppFile S1**.

### DNA extraction and qPCR assays

DNA extraction and qPCR assays were performed as in (Brunet *et al*., 2023). Briefly, environmental DNA from swabs was extracted using the DNA/RNA Miniprep kit (ZymoBiomics) following the manufacturer’s instructions. DNA was quantified with the QuantiFluor dsDNA System (Promega) kit and samples normalized at 0.5 ng.μl^-1^ before qPCR and library preparation. DNA concentrations obtained from each sample are listed in **SuppFile S1**.

The number of total 16S rRNA gene copies was assessed using qPCR (primers 926F: 5’- AAACTCAAAKGAATTGACGG-3’ /1062R: 5’-CTCACRRCACGAGCTGAC-3’, (Bacchetti De Gregoris *et al*., 2011) on a LightCycler 480 Instrument II (Roche). The global predicted coverage of this primer pair is 94.1% of all bacterial sequences present in Silva SSU r138.1 (Silva Testprime with only one mismatch allowed, analysis performed in June 2024). All bacterial phylum-level taxa in the Silva database had predicted coverages above 90%, except 10-bav-F6 (33%), *Candidatus* Bipolaricaulota (*Acetothermia* 50%), *Bdellovibrionota* (85%), *Chrysiogenetota* (75%), *Candidatus* Dependentiae (88%), *Dictyoglomota* (88%), *Candidatus* Edwardsbacteria (75%), NKB15 (86%) and *Patescibacteria* (46%). Among these, only *Bdellovibrionota*, *Candidatus* Dependentiae and *Patescibacteria* were detected in our 16S rRNA gene metabarcoding dataset, with average relative abundance of 1.2, 0.0001 and 0.09%, respectively. qPCR results are listed in **SuppFile S1,** and MIQE information related to qPCR experiments is available in **SuppFile S2**.

### Library preparation, sequencing and sequence processing

The V3-V4 region of 16S rRNA gene was amplified using the primers S-D-Bact-0341-b-S-17 (5’ CCTACGGGNGGCWGCAG 3’) and 799F_rc (5’ CMGGGTATCTAATCCKGTT 3’ (Thomas *et al*., 2020). Sequencing was conducted on a MiSeq paired-end sequencing run (300 cycles × 2, Illumina, San Diego, CA, USA) as described previously (Thomas *et al*., 2020). 16S rRNA gene amplicon sequences are available at NCBI under BioProject PRJNA1135191.The R package *DADA2* (version 1.22.0, Callahan et al. 2016) was used to exclude primers sequences, filter and trim low quality sequences (truncLen=c(280,210), trimLeft = c(10,10), minLen = 150, maxN=0, maxEE=c(2,3), truncQ=2), for denoising, paired-read merging, inferring Amplicon Sequence Variants (ASVs, length between and 400 and 452 nucleotides), chimera removal and taxonomic assignment using the SILVA 138.1 (Quast *et al*., 2013; Yilmaz *et al*., 2014) reference database. Default parameters were used unless otherwise specified. All sequences affiliated to 16S rRNA gene from chloroplasts, mitochondria, cyanobacteria, Eukaryota and *Archaea*, as well as sequences found in negative controls (PCR molecular grade water) were removed using the R package *phyloseq* (version 1.38.0, McMurdie and Holmes, 2013).

### Multivariate and statistical analyses

Relative ASV abundance was multiplied by the total number of 16S rRNA gene copies quantified with qPCR for each sample, to obtain a table of abundance of each ASV that was used in all subsequent analyses. To assess dissimilarities in community structure between algal tissues, Bray–Curtis dissimilarity index (Bray *et al*., 1957) was calculated before non-metric multidimensional scaling (NMDS). Permutational analysis of variance (PERMANOVA, 999 permutations) was calculated using the adonis function to discriminate groups of samples according to algal tissues or seasons (Anderson, 2001), followed by multivariate pairwise comparisons using pairwise.perm.manova. Mean dispersions within groups were calculated using betadisper (999 permutations). To identify environmental parameters that may be associated with microbial community structure, we performed distance-based redundancy analysis (dbRDA; Legendre and Anderson, 1999) on the Bray–Curtis dissimilarity matrix with dbrda function. To avoid having highly correlated variables, pH, SiOH4, NOP, PO4, NO3, NH4, DN15 were removed prior to the analysis. Only significant environmental factors were selected for dbRDA visualization (ANOVA, P < 0.05). Months were grouped in 3 different seasons (Winter: Feb, Mar, Jan; Summer: Jun, Jul, Aug; Autumn: Sep, Oct, Nov, Dec) prior to PERMANOVA and betadisper analysis. Differential sequence abundance analysis between the 3 seasons (qPCR-corrected counts from months of the same season were averaged) was performed at the ASV level using the GLM test with the ALDEx2 R package (Fernandes *et al*., 2014). To detect differential abundance, this test uses ratios among taxa, which are conserved regardless of whether data are relative or absolute.

A standardized approach based on occupancy–redundancy distribution was applied to identify core genera on each algal tissue (Shade and Stopnisek, 2019). Using a presence/absence matrix (1 for presence, 0 for absence), an index was calculated for each genus, as the sum of the occupancy term and the redundancy term for each month. The occupancy term is the replicate average of the presence/absence matrix. The redundancy term is equal to 1 if the genus is present in all three replicates and to 0 otherwise. The index was scaled to an upper limit of 1 and calculated separately for each algal tissue.

### Phylogenetic analysis

Phylogenetic analysis was conducted using phylogeny.fr (Dereeper *et al*., 2008) for the most abundant *Granulosicoccus*-affiliated ASVs in the dataset (representing at least 5% of the *Granulosicoccus* community in at least 5 samples) together with 16S rRNA gene sequences from cultured *Granulosicoccus* strains and uncultured clones retrieved from Genbank. Sequences were aligned using Muscle (full mode) and the resulting alignment was curated using Gblocks. The 405 conserved positions were used for neighbor-joining tree reconstruction with K2P substitution model and 1,000 bootstraps. The tree was visualized using iTOL (Letunic and Bork, 2024) and rooted at midpoint.

## Results

The structure and dynamic of the bacterial community associated with five phylogenetically distant macroalgae from a temperate rocky shore has been investigated over one year. The ASV relative abundance was multiplied by the total number of 16S rRNA gene copies to consider absolute taxa count and obtain a more precise understanding of the algal epimicrobiome structure.

### Algal host as a major driver of epiphytic bacterial community composition

The quantified number of bacterial 16S rRNA gene copies (**SuppFile S1**) varied significantly along the year (2-way ANOVA, ‘sampling month’ effect F_9,158_ = 5.0, p < 0.001) but not across macroalgal species (F_4,158_ = 0.8, p = 0.531) and with no interactions between the two factors (F_36,158_ = 1.1, p = 0.3). This proxy for total bacterial abundance was relatively homogeneous for all algae, with year-round average ranging between 7 x 10^6^ to 1 x 10^7^ 16S rRNA gene copies.cm^-2^, depending on the algal species. Yet, we could detect a large range of variations spanning more than two orders of magnitude, from a minimum of 1.04 x 10^5^ (Ldig-base in February 2020) to a maximum of 5.02 x 10^7^ 16S rRNA gene copies.cm^-2^ (Ldig-old in November 2020). The structure of the epiphytic bacterial communities was strongly impacted by the algal host (**Figure 1A**; PERMANOVA, F_4,203_ = 22.7, P < 0.001), showing that communities from different algal species sharing the same habitat are distinct from one another. This observation is supported by the fact that 80% of the total number of ASVs (8 316 out of 10 243) are specific to an algal species (**SuppFigure S1**) and only 1% (103) are present on all algae. However, 83% of these specific ASVs were only found in one sample. Despite their closer phylogenetic distance, brown algae (Ldig, Anod and Fser, all belonging to the class Phaeophyceae) did not share more ASVs or genera than with green (Ulva) and red (Ppal) algae. Besides host taxonomy, the age of algal tissue also influenced the surface density and composition of the microbiota. Indeed, the younger meristematic tissues of *L. digitata* (Ldig-base) consistently harbored less 16S rRNA gene copies than the medium frond (Ldig-medium, paired t-test, P = 0.002) and the older apical tissues (Ldig-old, paired t-test, P < 0.001). Multivariate analysis also separated Ldig-base epibacterial communities from Ldig-medium and Ldig-old (**Figure 1B**; PERMANOVA, F_2,86_ = 12.4, P < 0.001).

**Figure 1:**
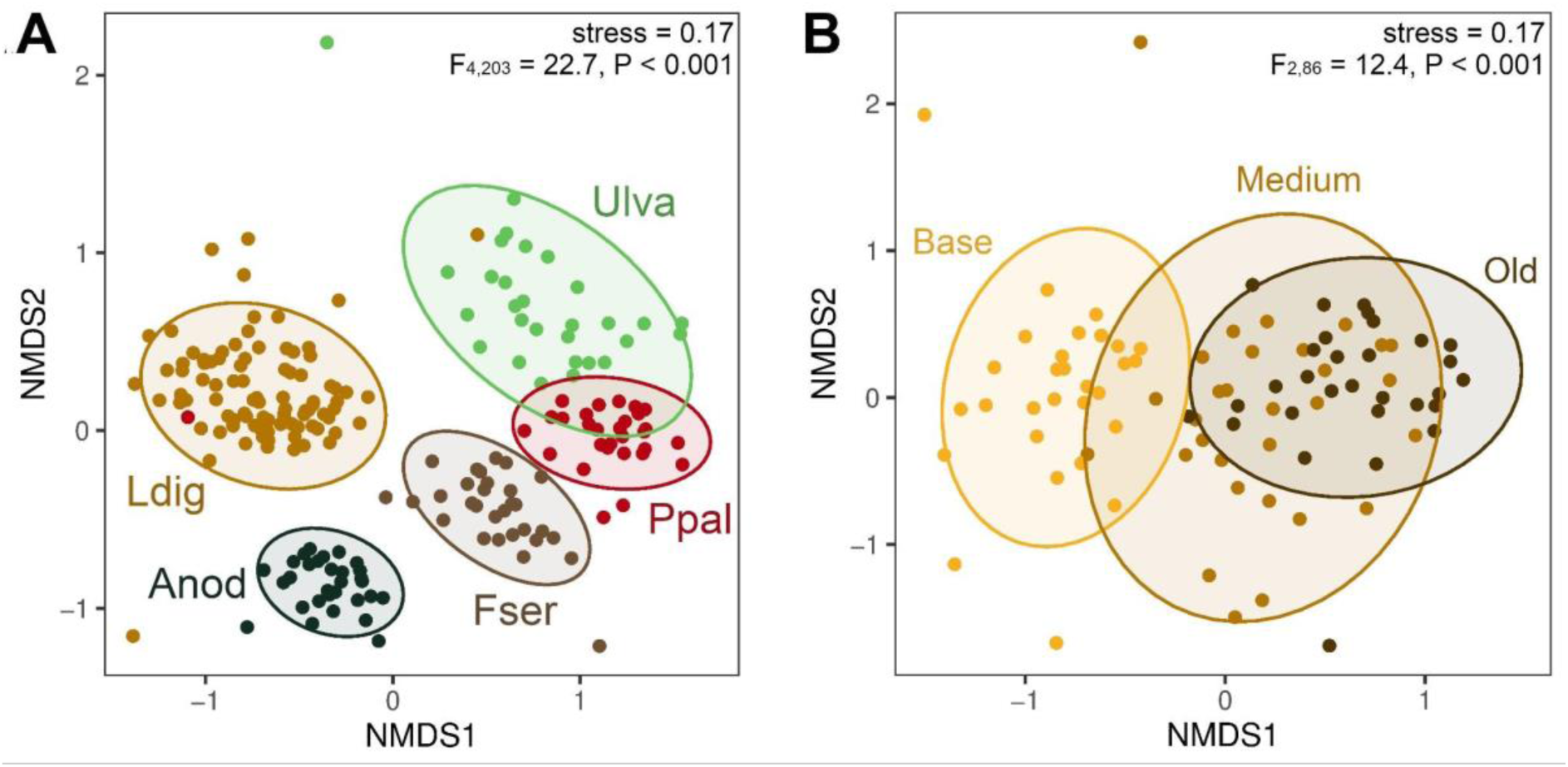
NMDS ordination plots of abundance data of all samples (**A**) or *L. digitata* samples (**B**), based on Bray-Curtis dissimilarity. 95% confidence ellipses for a multivariate t-distribution are depicted for the different type of algal tissues (either algal species or *L. digitata* blade part). PERMANOVA F-statistic and p-value are displayed.

*Gammaproteobacteria*, *Alphaproteobacteria*, *Bacteroidia, Acidimicrobiia and Planctomycetes* were the most abundant bacterial classes on all algae, representing on average 93-98 % of the communities, depending on the algal host (**SuppFigure S2**). After pooling the ASVs by genus, we observed that 28% of all genera (93 out of 331) were shared between all algae, suggesting they might be ubiquitous algal epibionts (**Figure 2** and **SuppFile S3**). These generalists were mainly *Bacteroidia* (including *Flavobacteriales*), *Gammaproteobacteria* and *Alphaproteobacteria* and represented on average 50-75 % of all sequences depending on algae. An occupancy-redundancy index was calculated following Shade and Stopnisek, 2019 method to identify core genera that were consistently found associated with the five studied algal species all year round (**Figure 2B**). Genera that had an index > 0.65 on all algae were defined as core genera: *Granulosicoccus* (*Gammaproteobacteria*)*, Litorimonas*, *Hellea*, *Fretibacter (Alphaproteobacteria)*, *Portibacter*, *Algitalea, Rubidimonas, Lewinella (Bacteroidota)*, *Blastopirellula* (*Planctomycetota*) and Sva0996 marine group (*Actinomycetota*). These represented only 3% of all genera, yet accounted for 46% of the bacterial abundance on average. The genus *Granulosicoccus* was the only one to have an index equal to 1 on all algal tissues, meaning it was present in all 209 samples (see below). On the contrary some genera were specifically associated with particular algae. Especially, the genus *Arenicella* (*Gammaproteobacteria*) was strongly associated with brown algae all year round (index > 0.78, 2.8 x 10^5^ copies.cm^-2^ on average) while scarce on Ppal (index = 0.27, 3.4 x 10^4^ copies.cm^-2^) and Ulva (index = 0.2, 4.1 x 10^3^ copies.cm^-2^). The opposite was observed for *Truepera* (*Deinococcota*; index < 0.3 on brown algae, > 0.9 on the red and green algae). Some genera were only found strongly associated (index > 0.8) with one algal species, namely *Tateyamaria* (*Alphaproteobacteria* on Anod, *Pibocella* (*Bacteroidota*) on Fser, *Sulfitobacter (Alphaproteobacteria)*, *Polaribacter (Bacteroidota)* and *Jannaschia* (*Alphaproteobacteria*) on Ulva. Moreover, important differences were observed between the different parts of the Ldig frond (**Figure 2**). In particular, members of the Sva0996 marine group and of the family *Microtrichaceae* (*Actinomycetota*) accounted on average for only 3 x 10^4^ 16S rRNA gene copies.cm^-2^ (1%) on the meristem compared to 2 x 10^6^ (15%) and 4 x 10^6^ (21%) on the medium and old frond respectively. These taxa were also overrepresented on Fser (2 x 10^6^, 21%) and Ppal (2 x 10^6^, 20%).

**Figure 2:**
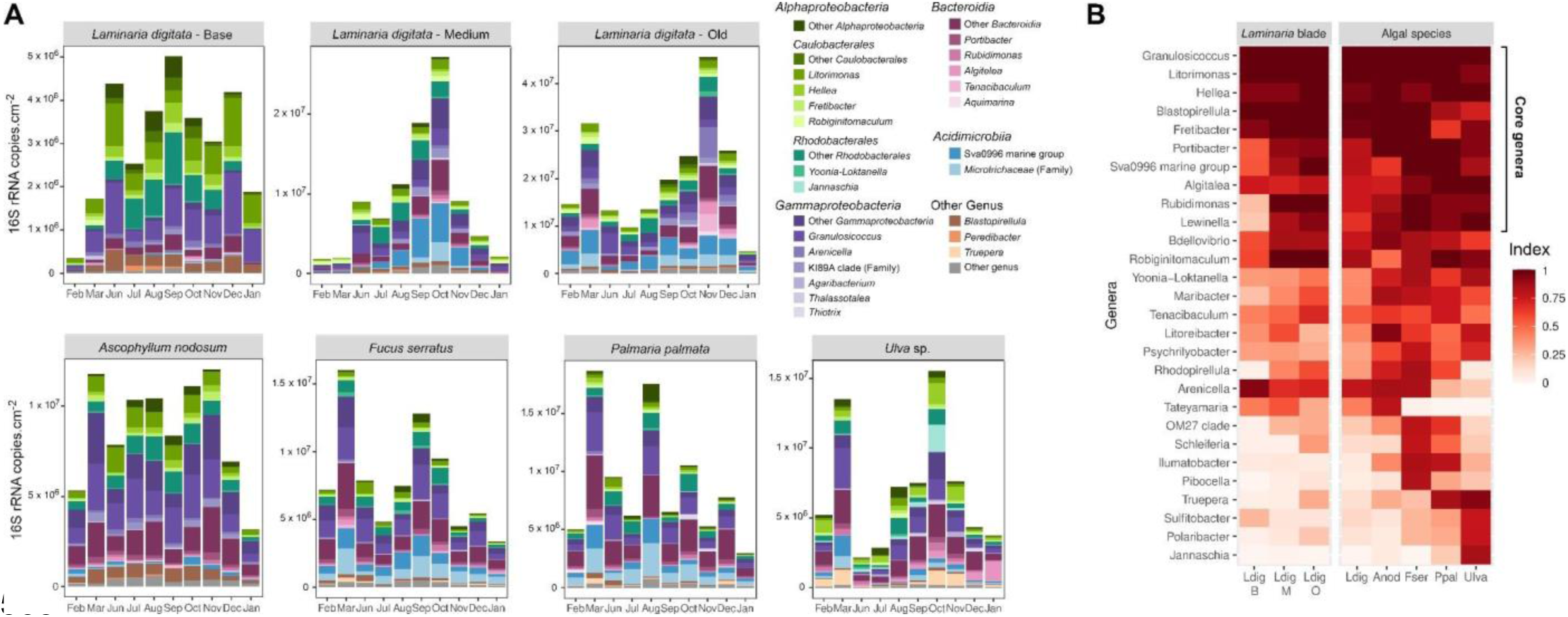
**(A)** Estimated abundance of the main bacterial genera. Only genera representing more than 5% (triplicate average) of the communities for at least one condition (ie. at one time point for one algae) are shown. **(B)** Approach to identify core macroalgal genera. Indices are calculated based on abundance-occupancy distributions (see Methods for index calculation). Only genera that have an index > 0.5 on at least one type of tissue are represented. For the Ldig column, the index is an average of the indices calculated on the base, medium and old Ldig tissues.

### Seasonal succession of epiphytic bacterial communities

The number of 16S rRNA gene copies.cm^-2^ varied differently through seasons depending on the algal host, and no common pattern to all algae could be highlighted (**Figure 3**). The number of total bacteria tended to reach a peak in late summer / autumn on Ldig, Fser, Ppal and Ulva. Ldig-old, Anod, Fser, Ppal and Ulva samples from March 2020 showed higher bacterial abundances than neighboring months (February and June). The absence of data in April and May (COVID lockdown) prevents us from assessing if this abundance peak was an isolated event caused by specific environmental conditions in March or if it was characteristic of through the entire spring season.

**Figure 3:**
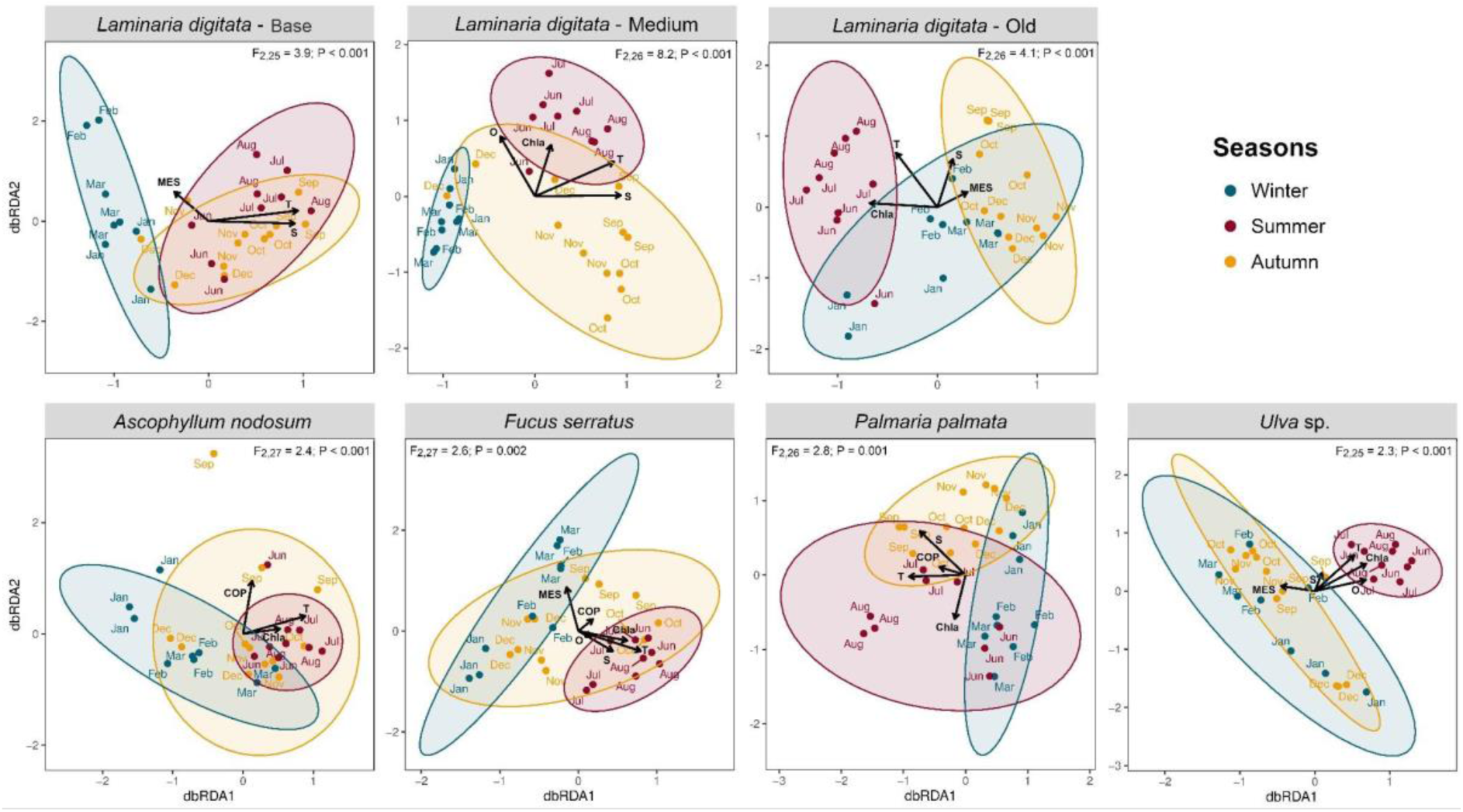
db-RDA plots with vectors representing significant contextual predictors. 95% confidence ellipses for a multivariate t-distribution are depicted for the different seasons. PERMANOVA F-statistic and p-value are displayed. T: Temperature; O: Dissolved oxygen; S: Salinity; ChlA: Chlorophyll A; MES: Suspended matter; COP: particulate organic carbon.

Globally, the number of observed ASVs did not significantly differ through the year (**SuppFigure S3**). A 3-fold decrease of the ASV number was only observed in June compared to October (Ldig-medium), December (Ldig-old) or September (Ulva). Distance-based ReDundancy Analysis (dbRDA) and PERMANOVA (**Figure 4**) showed clear dissimilarities between the three identified seasons (winter, summer and autumn) for all algae (except between winter and autumn with Anod and Ulva) and suggested these seasonal variations might be partly explained by environmental parameters, especially seawater temperature, salinity and chlorophyll A concentration.

**Figure 4:**
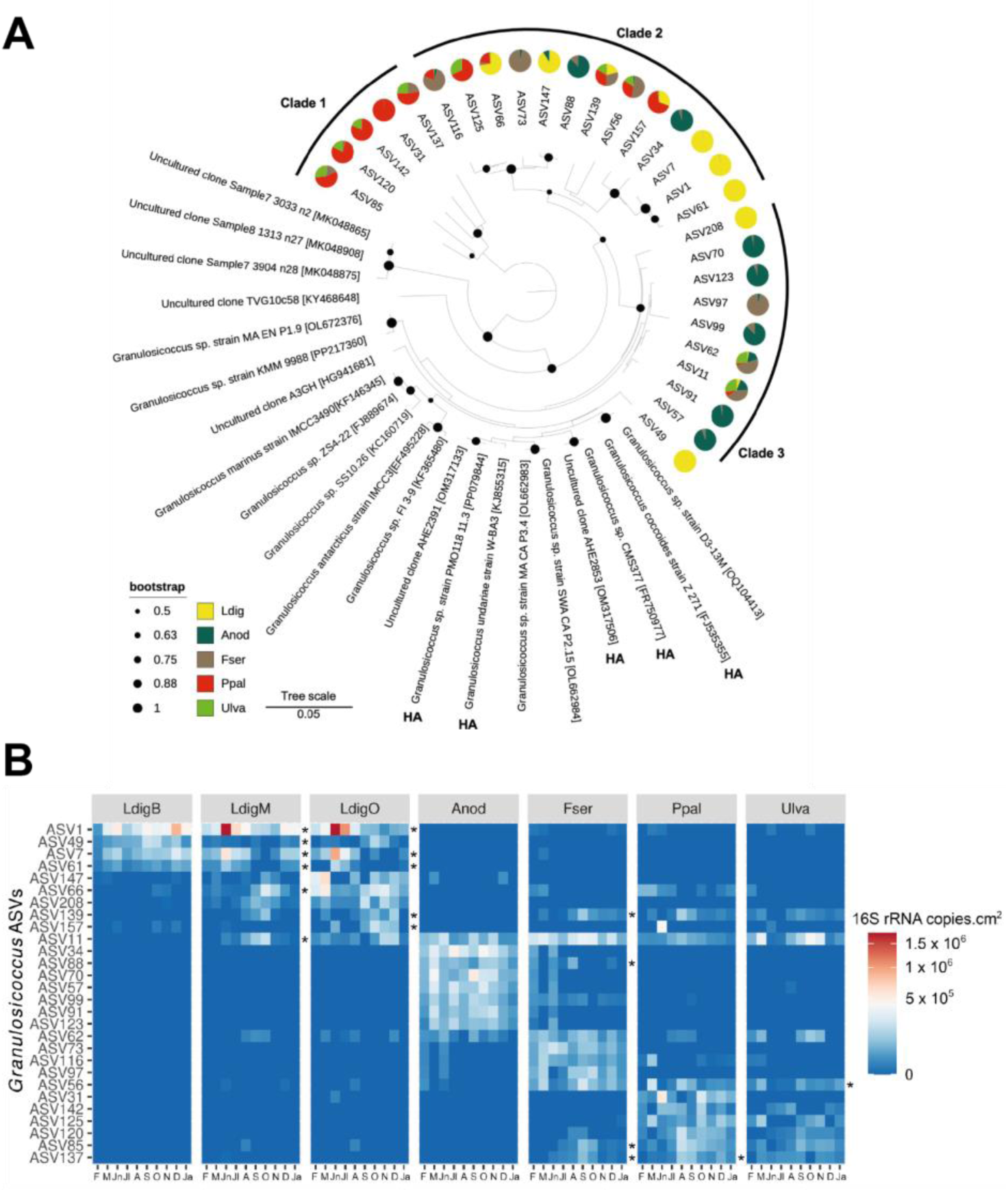
**(A)** Phylogenetic tree of the main 16S rRNA ASVs affiliated to *Granulosicoccus* and reference sequences (Genbank accession in brackets). Pie charts represent year-round average abundance of each ASV on different algal species. HA: host-associated. **(B)** Heatmap of the abundance of the main ASVs assigned to the genus *Granulosicoccus*. Only *Granulosicoccus* ASVs representing at least 5% of the *Granulosicoccus* community in at least 5 conditions (out of 70) are displayed. Abundances of triplicates were averaged and square root transformation was applied. Asterisks on the right panel of each alga denote differentially abundant ASVs through seasons. F: February; M: March; Jn: June; Jl: July; A: August; S: September; O: October; N: November; D: December; Ja: January.

No single ASV was found to vary consistently through seasons on all algae (**SuppFile S4**). This might be partly explained by the low proportion of ASVs present on all algae, as mentioned above. Anod-associated communities were the least impacted by seasons with only 9 ASVs differentially abundant (**SuppFile S4, Figure 2A**). Bacterial communities’ seasonal dynamics varied depending on the part of the Ldig blade. The basal part exhibited less variations than the older ones (15 differentially abundant ASVs compared to 55 and 92 on medium and old fronds, respectively). On the basal tissues, 2 ASVs from the genus *Peredibacter* (*Bdellovibrionota*) were more abundant during summer months (3 x 10^4^ 16S rRNA gene copies.cm^-2^, against 2 x 10^2^ in winter and 6 x 10^3^ in autumn, respectively) and 3 ASVs from the family *Rhodobacteraceae* (*Alphaproteobacteria*) less abundant in winter. The most outstanding abundance variations on the older tissues were observed for ASVs from the classes *Acidimicrobiia* (Sva0996 marine group and the family *Microtrichaceae*) and *Bacteroidia* (orders *Chitinophagales* and *Flavobacteriales*), that peaked in autumn on the medium frond and in autumn and winter at the tip part. The community succession on Fser was characterized by a significantly higher abundance of *Micavibrionales* (*Alphaproteobacteria*) in summer (2 x 10^4^ 16S rRNA gene copies.cm^-2^, against 0 in winter and 2 x 10^3^ in autumn). Also, the abundance of 28 *Chitinophagales* (*Bacteroidota*) ASVs significantly varied between seasons but exhibited distinct abundance patterns. Only few variations were observed with the red alga Ppal (24 differentially abundant ASVs). The most notable succession was a peak of the *Aquimarina* (*Flavobacteriales*) population in winter (1 x 10^5^ 16S rRNA gene copies.cm^-2^ compared to none in summer and 4 x 10^3^ autumn). Summer communities displayed high specificity at the surface of Ulva. Thirty-one ASVs were differentially abundant, in particular, members of the genus *Truepera* (*Deinococcales*), of the families *Micotrichaceae* (*Actinomycetota*) and *Saprospiraceae* (*Bacteroidota*) exhibited a sharp decrease in summer while alphaproteobacteria from orders *Rhodobacterales* and *Caulobacterales* (*Litorimonas* and *Fretibacter*) observed an opposite trend.

### *Granulosicoccus* are ubiquitous members of macroalgal epibiota

The genus *Granulosicoccus* (*Gammaproteobacteria*) was the only one present in all 208 samples, covering the 5 different algal species and the 10 sampling months. It was also the most abundant genus, representing 15.4% of the communities on average (up to 42% on Ldig-base), and accounting for 1.2 x 10^6^ copies.cm^-2^ on average (up to 7.7 x 10^6^ copies.cm^-2^ on Ppal). It comprised 464 different ASVs, the highest number of ASVs for all genera in our dataset, suggesting a large micro-diversity. Half of these ASVs (236) were only found in one condition (ie. on one type of tissue at one specific time point). Phylogenetic analysis of the most abundant *Granulosicoccus* ASVs (representing at least 5% of the *Granulosicoccus* community in at least 5 conditions) confirmed this large diversity and separated three main clades (**Figure 4**), likely representing new species compared to the 4 validly described species. Clades 2 and 3 clustered with reference sequences from cultured *Granulosicoccus* strains and uncultured clones. This notably included other marine host-associated strains, retrieved from the brown alga *Undaria*, the red alga *Gracilaria*, the seagrass *Zostera marina*, sponge and plankton (Kurilenko *et al*., 2010; Park *et al*., 2014; Heins and Harder, 2023). Clade 1 comprised 5 ASVs that clustered away from other sequences. In addition, all ASVs were not equally distributed on the different algal species. Only two closely related *Granulosicoccus* ASVs were consistently detected on all algal species (ASV11 and to a lesser extent ASV62). Most of the ASVs were specific or strongly linked to a unique algal species, e.g. ASV1, ASV7, ASV61, ASV208 and ASV49 to Ldig, ASV34, ASV70, ASV123 ASV91 and ASV57 to Anod, ASV73 to Fser and ASV31 to Ppal (**Figure 4A**). While clade 2 encompassed ASVs detected on multiple algae, clade 1 and clade 3 comprised ASVs more abundant on red and brown algae, respectively. *Granulosicoccus* ASVs associated with medium and old Ldig tissues exhibited stronger significant seasonal variations than *Granulosicoccus* ASVs associated with the other algal species (**Figure 4B**). Moreover, even ASVs found on the same tissue displayed distinct variations patterns, e.g. for Ldig-medium where ASV1, 7 and 61 peaked in summer and ASV66, 208 and 139 in autumn.

## Discussion

Next-generation sequencing methods have become indispensable tools in microbial ecology, opening new avenues for studying microbial communities. However, these approaches are limited, as they yield relative abundance profiles of microbial taxa, which may not accurately reflect true variations in the actual taxon abundances in the environment (Gloor *et al*., 2017; Lloréns-Rico *et al*., 2021). In this study, we estimated actual ASV abundance by multiplying the ASV relative abundance matrix by the corresponding number of 16S rRNA gene copies estimated through qPCR (Barlow *et al*., 2020; Jian *et al*., 2020). This approach has been previously applied to soil microbiota (Lou *et al*., 2018; Azarbad *et al*., 2022), revealing that quantification provides a more comprehensive understanding of bacterial community dynamics than compositional data, which are more prone to false-positive changes. Absolute abundance of bacterial taxa associated with macroalgae has been previously characterized but focusing only on a few targeted taxa using specific qPCR primers or FISH probes (Ramírez-Puebla *et al*., 2022; Brunet *et al*., 2023). Here we implemented this method to quantitatively characterize the whole macroalgal epimicrobiota composition for the first time, aiming to assess true taxa variations across different algal hosts and seasons. Although providing valuable improvements over sole amplicon sequencing to estimate absolute abundance of different taxa, coupling with qPCR still has some limitations. At least three limitations are common with the metabarcoding- only approach: (1) potential saturation of the swabs during sampling that could lead to underestimate bacterial abundance on densely colonized algal surfaces; (2) variation in extraction efficiencies depending on algal tissue composition (e.g. polysaccharide content); (3) unequal lysis of different bacterial taxa. Here, we used flocked nylon swabs, previously shown to outperform other swab types in terms of collection efficiency and DNA recovery from microbiomes (Bruijns *et al*., 2018; Wise *et al*., 2021). The large variation in DNA concentrations retrieved from swabs (minimum 0.11 ng.µl^-1^, median 7.57 ng.µl^-1^, maximum 41.46 ng.µl^-1^, 92% of samples below 24 ng.µl^-1^, **SuppFile S1**) further suggests that if saturation occurred, it only affected a limited number of samples. Moreover, DNA recovery from swabs did not differ depending on algal species (on average Anod: 9.3 ng.µl^-1^; Fser 8.9 ng.µl^-1^; Ldig 10.3 ng.µl^-1^; Ppal 10.6 ng.µl^-1^; Ulva 6.9 ng.µl^-1^, **SuppFile S1**), suggesting tissue composition did not largely affect extraction efficiency. An additional bias lies in the use of different sets of primers for qPCR and amplicon sequencing. Indeed, qPCR requires short amplicon size for optimal efficiency and specificity (< 200 bp, Debode et al., 2017), whereas optimal primers for 16S metabarcoding cover a region of at least approximately 450 bp after merging, to encompass both conserved and hypervariable regions. Yet, this bias is minimal since the qPCR primers covered most of the taxa found in the 16S metabarcoding dataset (see Methods).

### Common core of the epiphytic microbiome

The large number of samples collected in this study, encompassing 7 types of algal tissues among 5 algal species at 10 sampling months, allow for the identification of a core epiphytic microbiome that would be consistently found on algae all year round. The identification of these core microbes is crucial as they likely have more substantial impact on the host’s biology than other members of the microbiome (Neu *et al*., 2021). Here the core microbiota was identified based on occupancy and redundancy criteria as recently used in soil microbial ecology studies (eg. Hodgson et al., 2024). The identified core genera, especially *Granulosicoccus*, *Litorimonas* and *Hellea*, concur with previous works that revealed the preponderance of these taxa at the surface of macroalgae (Weigel and Pfister, 2019; Lemay *et al*., 2021; Wood *et al*., 2022; Burgunter-Delamare *et al*., 2023; Lu *et al*., 2023). *Granulosicoccus* was the dominant genus on all algae, both in terms of presence/absence and abundance. Predominance of *Granulosicoccus* on the surface of other brown algae was previously suggested using relative compositional data (eg. Lemay et al., 2021; Weigel et al., 2022; Burgunter-Delamare et al., 2023) and quantitative cell counts using FISH probes (Ramírez-Puebla *et al*., 2022). Here we quantify this ubiquity on a larger temporal scale and on other algal species, extending this observation to red and green algae. Using an estimate of 3 rRNA operons per cell (as seen in the complete reference genome of *Granulosicoccus antarcticus* (Kang *et al*., 2018)), the maximum observed density of 7.7 x 10^6^ 16S rRNA gene copies.cm^-2^ on the red algae *P. palmata* would correspond to ca. 2 millions *Granulosicoccus* cell.cm^-2^. Using FISH counts on the kelp *Nereocystis luetkeana* in summer, Ramírez-Puebla et al., 2022 reported 5.5 x 10^4^ and 6.5 x 10^6^ *Granulosicoccus* cells.cm^-2^ on basal and old tissues, respectively. Comparable values can be inferred from our dataset on the basal and old tissues of the kelp *L. digitata* in summer (5 x 10^5^ and 2.3 x 10^6^ *Granulosicoccus* cells.cm^-2^, respectively), confirming qPCR correction of metabarcoding compositional data is a useful approach to estimate absolute abundances of specific taxa. *Granulosicoccus* sequences were scattered into multiple ASVs that fall within distinct clades whose abundance patterns across algal hosts and seasons differ. These clades exhibit distinct tissue specificity and could be characterized either as generalist (clade 2) for containing taxa colonizing all types of tissue or specialist for clades 1 and 3 that encompass taxa that are more specific to particular hosts. These data indicate that ASVs from a same genus colonize distinct ecological niches and underscore important genetic and functional diversity within core bacterial genera associated with macroalgae. Similar observations were made in recent studies where different genomes of core epiphytic genera, especially *Granulosicoccus*, were differentially abundant at the surface of the bull kelp *N. luetkeana* depending on its location (Weigel *et al*., 2022) or on *Fucus* sp. depending on season and geographical location (Park *et al*., 2022). The analysis of *Granulosicoccus* genomes revealed strong abilities for an epiphytic lifestyle, that would allow fast colonization and stable associations with the host. In particular, genes involved in motility and chemotaxis, B12 vitamin biosynthesis, DMSP metabolism, transport and utilization of sugars, were found abundant in *Granulosicoccus* metagenomes (Kang *et al*., 2018; Weigel *et al*., 2022).

### Host algal tissue is a major driver of epiphytic community composition

Our study revealed that the type of algal tissue, whether it is different algal species or different blade parts, was a stronger driver of epiphytic community composition than seasonality. This observation is consistent with previous reports stating that host characteristics may have a greater impact on shaping the structure of macroalgae-associated microbial communities than environmental variables (Marzinelli *et al*., 2015; Burgunter-Delamare *et al*., 2023). Chemical composition of algal tissues varies depending on algal taxonomy (Mišurcová, 2011) or tissue age (Küpper *et al*., 1998). Significant differences of concentration of nutritional factors such as proteins, minerals, lipids or sugars were largely shown. Moreover, polyphenol content was reported to be 10-20 times higher in *A. nodosum* and *F. serratus* compared to the other algae collected in this study (Pandey *et al*., 2022). Species-specific differences were also reported for iodine, as its concentration was shown to be approximately 5 and 20 times higher in *L. digitata* compared to *A. nodosum* or *F. serratus* and *P. palmata* or *Ulva* sp., respectively (Nitschke *et al*., 2018). Both iodine and polyphenols are known to be algal defense compounds involved in the regulation of microbial colonization (eg. Cosse et al., 2009; Besednova et al., 2020). Such chemical variations were also observed across seasons but were much less pronounced than between algal species (Nitschke *et al*., 2018; Pandey *et al*., 2022). Therefore, if these compounds are drivers of community composition, their limited annual variations within the tissues could partly explain why seasons had a smaller impact on community composition than algal host. Interestingly, microbiota composition similarities were not higher between brown algae (*L. digitata*, *A. nodosum* and *F. serratus*) than with the red or green algae. Similarly, no common pattern was found among species inhabiting the same intertidal zone. Then, despite the important discrepancies observed in the microbiota of distinct algal species, neither higher taxonomic level (algal phylum) nor algae location on the foreshore appeared to be strong community drivers, corroborating observations made by Chen et al., 2022 and Selvarajan et al., 2019. Nevertheless, the abundance of certain taxa displayed important differences between algal phyla, *Arenicella* was consistently associated only with brown algae, whereas *Truepera* was consistently found only with red and green algae. *Truepera* phylum specificity was also reported in (Lu *et al*., 2023) with different algal species.

This work shows that different parts of the *L. digitata* blade support distinct bacterial communities and that total bacteria abundance and diversity are higher on older tissues, in accordance with studies carried out on the same species (Ihua *et al*., 2020) and on other kelp species (Bengtsson *et al*., 2012; Weigel and Pfister, 2019; Lemay *et al*., 2021; Ramírez-Puebla *et al*., 2022; Burgunter-Delamare *et al*., 2023). Moreover, the specific colonization of older *L. digitata* tissues by members of the Sva0996 marine group, in comparison to the meristem, concur with the one observed in (Brunet *et al*., 2021) where the abundance of this taxon was linked with tissue decomposition and degradation. Then, the presence of such taxa on old damaged tissues might be related to their capacity to use complex algal derived organic matter such as polysaccharides (Brunet *et al*., 2022).

Seasonal variations of the surface microbiota have received unequal attention depending on the algal host. To our knowledge, to date, there is no report of these fluctuations on *Ascophyllum* sp. or *Palmaria* sp., while a few studies were conducted on *Fucus* sp. (Lachnit *et al*., 2011; Park *et al*., 2022), *Ulva* sp. (Tujula *et al*., 2010; Lachnit *et al*., 2011) and *Laminaria* sp. (Corre and Prieur, 1990; Bengtsson *et al*., 2010; Brunet *et al*., 2021). Seasonal microbial variations are known to be influenced by a combination of biotic and abiotic factors. Here we showed that seasons affect epiphytic community composition for all algae but in different ways. Community richness decreased in summer for *L. digitata* and *Ulva* sp. Temperature and salinity were environmental parameters that could explain the observed seasonal discrepancies. Seawater temperature shapes microbial community variations, in particular, it was already shown that increasing temperatures lead to a decrease in community richness on kelp (Paix *et al*., 2021) and in seawater (Sunagawa *et al*.). Salinity was also reported as an important structuring variable of *Ulva*-associated bacterial communities (van der Loos *et al*., 2023). Anod-associated microbiota was the most stable across seasons among all five algae. *A. nodosum* is known to observe monthly epidermal shedding, a way for longer-lived marine macroalgae to rid themselves of epibionts via the removal of the outer cell layer of the thallus (Halat *et al*., 2015; Garbary *et al*., 2017). Therefore, it is likely that the newly exposed thallus, free of epibionts, does not present any alterations or damages and is monthly colonized by the same early bacterial epibionts.

This study, through the extensive number of collected samples and the analysis of absolute taxa abundance, revealed true microbiota variations based on hosts and seasons, providing novel insights in the structure and fluctuations of bacterial communities associated with macroalgae. Future studies should focus on deciphering the ecological and metabolic functions of the identified core epiphytic taxa to gain further insights into epiphyte–host interactions.

## Supporting information

SuppFile S1

SuppFile S2

SuppFile S3

SuppFile S4

## Acknowledgments

The authors thank Manon Choulot for her help during sample collection, and Dr. Simon Dittami for useful discussions on the manuscript. This work has benefited from the facilities of the Genomer platform and from the computational resources of the ABiMS bioinformatics platform (FR 2424, CNRS-Sorbonne Université, Roscoff), which are part of the Biogenouest core facility network. This work was funded by the French Government via the National Research Agency programs ALGAVOR (ANR-18-CE02-0001-01) and IDEALG (ANR-10-BTBR-04).

## Contributions (following CRediT taxonomy)

Conceptualization, Funding acquisition, Project administration: FT; Data curation, Formal analysis, Visualization, Writing-Original draft preparation: MB, FT; Investigation: MB, NLD, FT; Supervision: FT, TB. Writing-Review & Editing: MB, NLD, TB, FT.

---

**Figure S1:**
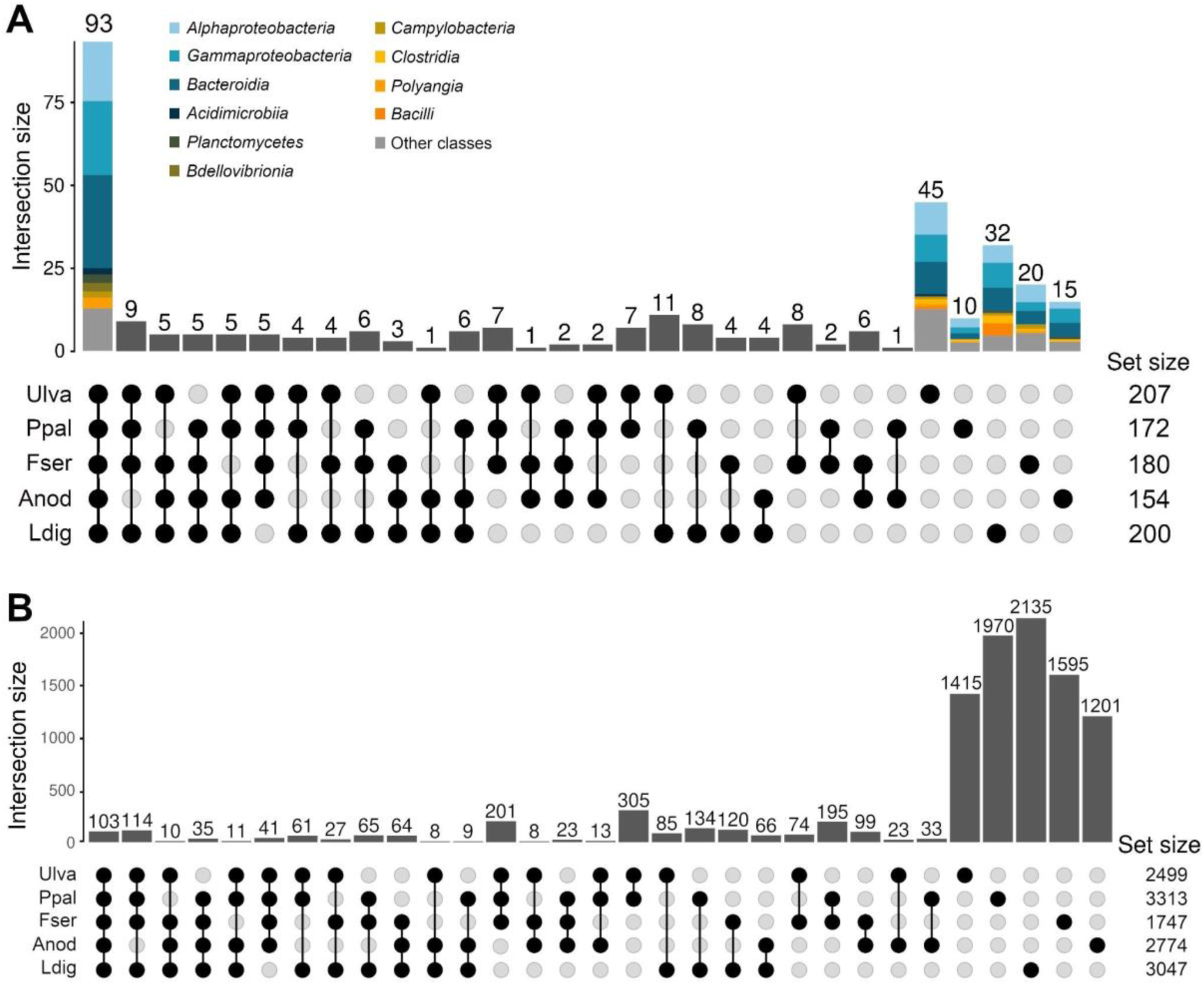
**(A)** Upset plot of the bacterial genera found in the algal microbiota. Set size represents the total amount of genera present on each alga. Taxonomy is displayed at the class level for the genera shared by all algae and the ones specific to one algal species. **(B)** Upset plot of the 10 243 ASVs found in the algal microbiota. Set size represents the total amount of ASVs present on each alga.

**Figure S2:**
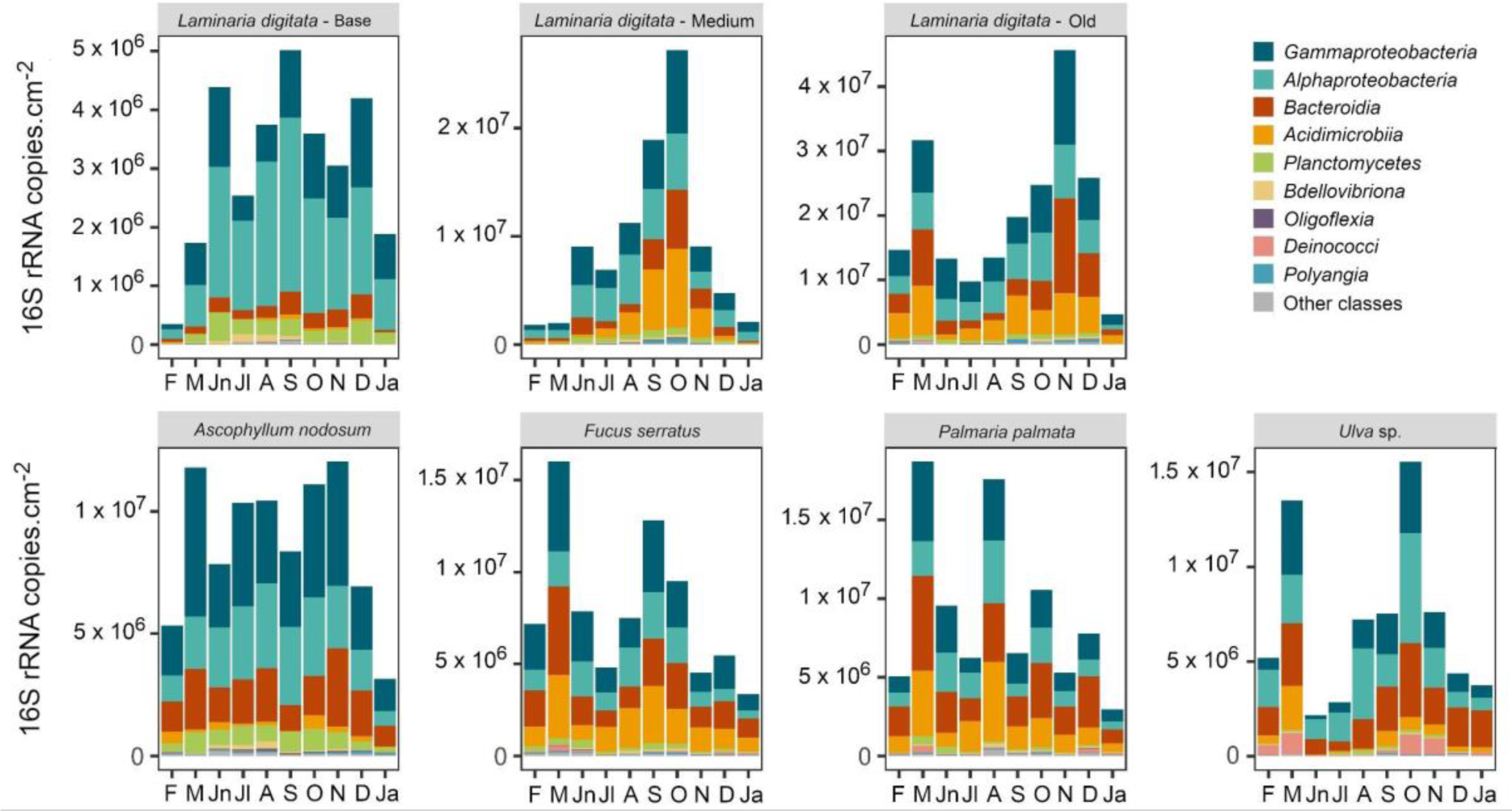
Estimated abundance of the main bacterial classes. Only classes representing more than 3% (triplicate average) of the communities for at least one condition (ie. at one time point for one algae) are shown.

**Figure S3:**
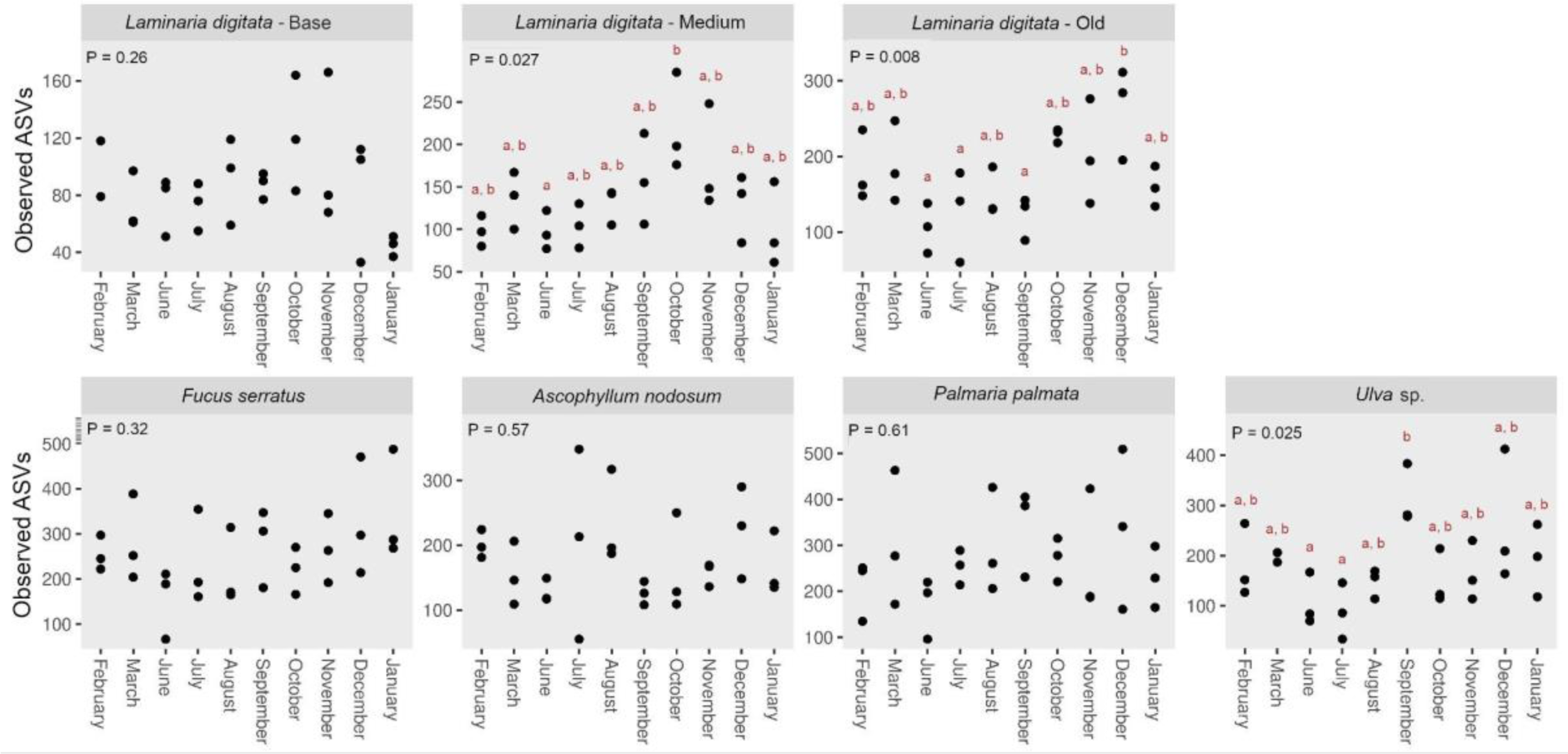
Fluctuation of the number of observed ASVs. When significant ANOVA results were found (P < 0.05), a post hoc Tukey HSD test was calculated. Accordingly, different letters indicate significant differences between sampling months.

**File S1:** Characteristic of the 209 samples of macroalgal epibiota collected during this study.

**File S2:** Details of the qPCR assay on the environmental samples, following the Minimum Information for publication of Quantitative real-time PCR Experiments (MIQE) guidelines.

**File S3:** List of the 331 assigned taxonomic genera. The number of samples in which they were found is indicated for each alga.

**File S4:** List of differentially abundant ASVs through seasons. A red excel cell indicates the abundance of the ASV varies significantly through seasons on the associated algal tissue. The different sheets named after each algal tissue display the season average abundance of the differentially abundant ASVs. The genus (or the deepest known taxonomic level) assigned to each ASV is mentioned.

